# Exaptation of two ancient immune proteins into a new dimeric pore-forming toxin in snails

**DOI:** 10.1101/2019.12.23.880021

**Authors:** M.L. Giglio, S. Ituarte, V. Milesi, M.S. Dreon, T.R. Brola, J. Caramelo, J.C.H. Ip, S. Maté, J.W. Qiu, L.H. Otero, H. Heras

**Affiliations:** Instituto de Investigaciones Bioquímicas de La Plata “Prof. Dr. Rodolfo R. Brenner”, INIBIOLP. CONICET CCT La Plata - Universidad Nacional de La Plata (UNLP), Facultad de Medicina,1900 La Plata, Argentina; Instituto de Estudios Inmunológicos y Fisiopatológicos, IIFP. CONICET CCT La Plata – UNLP, Facultad de Ciencias Exactas, 1900 La Plata, Argentina; Instituto de Investigaciones Bioquímicas de Buenos Aires, IIBBA. CONICET- Fundación Instituto Leloir, Av. Patricias Argentinas 435, C1405BWE Buenos Aires, Argentina; Department of Biology, Hong Kong Baptist University, 224 Waterloo Road, Hong Kong, China; Plataforma Argentina de Biología Estructural y Metabolómica PLABEM, Av. Patricias Argentinas 435, C1405BWE, Buenos Aires, Argentina

**Keywords:** Pore-forming toxin, poisonous snail egg, PmPV2, lectin, AB toxin, *Pomacea*, chemical defense, negative stain electron microscopy, small-angle X-ray scattering

## Abstract

The Membrane Attack Complex-Perforin (MACPF) family is ubiquitously found in all kingdoms. They have diverse cellular roles but MACPF but pore-forming toxic function are very rare in animals. Here we present the structure of PmPV2, a MACPF toxin from the poisonous apple snail eggs, that can affect the digestive and nervous systems of potential predators. We report the three-dimensional structure of PmPV2, at 15 Å resolution determined by negative stain electron microscopy (NS-EM) and its solution structure by small angle X-ray scattering (SAXS). We found that PV2s differ from nearly all MACPFs in two respects: it is a dimer in solution and protomers combine two immune proteins into an AB toxin. MACPF chain is linked by a single disulfide bond to a tachylectin chain, and two heterodimers are arranged head-to-tail by non-covalent forces in the native protein. MACPF domain is fused with a putative new Ct-accessory domain exclusive to invertebrates. Tachylectin is a six-bladed β-propeller, similar to animal tectonins. We experimentally validated the predicted functions of both subunits and demonstrated for the first time that PV2s are true pore-forming toxins. The tachylectin ^..^B^..^ delivery subunit would bind to target membranes, and then its MACPF ^..^A^..^ toxic subunit disrupt lipid bilayers forming large pores altering the plasma membrane conductance. These results indicate that PV2s toxicity evolved by linking two immune proteins where their combined preexisting functions give rise to a new toxic entity with a novel role in defense against predation. This structure is an unparalleled example of protein exaptation.

## Introduction

The integrity of cellular membranes is crucial for life and the disruption of such integrity causes cell death. Animals have evolved many strategies for damaging membranes and pore formation by proteins is frequently used in toxic attack on cells, as it can lead to efficient disruption of cell metabolism or even cell death. Among these proteins, the largest group belongs to the Membrane Attack Complex and Perforin / Cholesterol-Dependent Cytolysins (MACPF/CDC) superfamily, with ubiquitous distribution in all kingdoms. Most characterized members of MACPF/CDC interact with membranes and form large pores (hence the name: pore-forming proteins, PFPs). MACPF proteins function in immunity and bacterial pathogenesis (Anderluh, Kisovec, Krasevec, & Gilbert, 2014), and a very small group, termed pore-forming toxins (PFTs), have a toxic function (Peraro & van der Goot, 2016). These PFTs are present in bacteria, protists and fungi, but are very rare in animals. In fact, MACPF PFTs were reported only in a vertebrate (stonefish) and two groups of invertebrates: Cnidaria (Anderluh et al., 2014; Ellisdon et al., 2015) and the apple snail *Pomacea canaliculata* (Dreon et al., 2013).

We focus on the PFTs from the poisonous eggs of *Pomacea* apple snails (Gastropoda: Ampullariidae). Among them, *Pomacea canaliculata* eggs contain the toxin perivitelin-2 (PcPV2), one of the most toxic egg proteins known (Heras et al., 2008). PcPV2 is composed of two subunits, a MACPF chain, and a tachylectin-like chain [member of the F-type lectin family (Bishnoi, Khatri, Subramanian, & Ramya, 2015)], termed PcPV2-67 and PcPV2-31, respectively (Dreon et al., 2013; Heras et al., 2008). Moreover, the egg fluid (PVF) of *Pomacea maculata*, a related species, also contains a PV2-67 and PV2-31 like proteins orthologous of the two PcPV2 subunits (Mu, Sun, Heras, Chu, & Qiu, 2017).

PcPV2 can be included into the AB toxins, a small group of toxic proteins found in bacteria (e.g. botulinum neurotoxins) and plants (e.g. Type-2 RIP), that play a role in pathogenic processes and embryo defense, respectively. AB toxins contain two moieties, the “A” moiety that modifies some cellular target leading to cell death and the “B” moiety, which has usually a carbohydrate binding module (CBM), that recognizes glycans of the cell membrane and acts as a delivery subunit (Odumosu, Nicholas, Yano, & Langridge, 2010). Some of the CBM properties of AB toxins have been recognized in PcPV2 as it can agglutinate erythrocytes and recognize intestinal cells (Dreon et al., 2013), however, little is known about its sugar specificity and toxic mechanism. PcPV2 is unique among AB toxins in that not only the B but also the A moiety - a MACPF - has also putative membrane binding capacity, although there is no experimental confirmation of the pore-formation capacity of PV2s. For instance, experiments with mice indicate that minute quantities of PcPV2 are lethal if they enter the bloodstream (Dreon et al., 2013; Heras et al., 2008). Eggs of *P. maculata* were also poisonous and caused lethal toxicity to mice by an unidentified factor. It was observed that after PVF inoculation, severe signs pointing to nervous disorders appeared while at longer periods, mice showed paralysis of the rear limbs and even death (Giglio, Ituarte, Pasquevich, & Heras, 2016). The poisonous eggs of *P. canaliculata* and *P. maculata*, have an additional line of defense advertising the noxiousness by a bright pink coloration that warns predators (aposematic coloration) (Fig. 1A). Apple snail defensive strategy pays off and as a result eggs have very few predators (Yusa, Sugiura, & Ichinose, 2000). Here, we identified PmPV2 as the toxic factor in *P. maculata* eggs and studied its structure and putative functions. We found that PmPV2 present unique structural features compared with other MACPFs, and demonstrated that both subunits – tachylectin and MACPF – are functional, being the first experimental validation of the pore-forming capacity of PV2 toxins.

**Figure 1.**
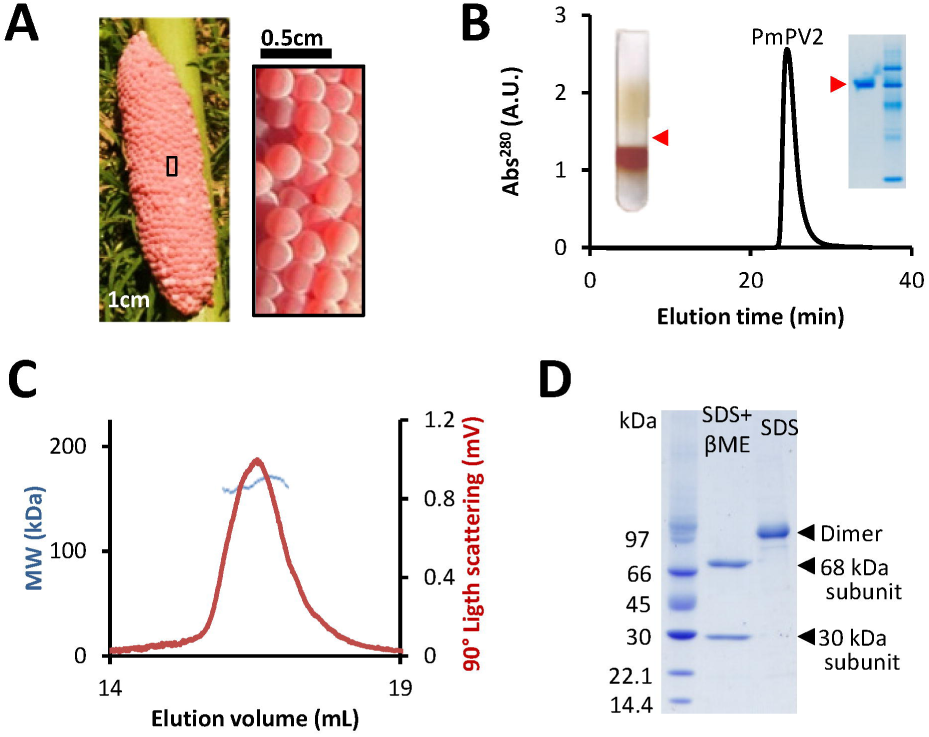
Identification of PV2 toxin from the poisonous eggs of *P. maculata*. (A) Egg clutch of the apple snail P. maculata. (B) Egg fluid from apple snail eggs was subjected to ultracentrifugation in NaBr gradient and isolation of native PmPV2 by ionic exchange and exclusion columns. *Insets:* ultracentrifugation tube showing PVF fractions and native-PAGE of purified PmPV2 (red arrowheads). (C) PmPV2 toxicity: Lethality of mice recorded after i.p. injection of PmPV2 and then fitted to a Hill equation (5 animals per group) (a); Cytotoxic effect on Caco-2 cells evaluated using MTT assay (b). (D) Molecular mass determination PmPV2 by SLS. (E) PmPV2 subunit composition analyzed by SDS-PAGE demonstrates the sample is dimeric with a single band corresponding to dimeric PmPV2 shown in lane SDS. Lane SDS+ βME shows a sample that has been deliberately monomerized following incubation with β-mercaptoethanol (βME) as reducing agent.

## Results

### Identification and toxic activity of PmPV2

We tracked the toxin of the PVF that causes lethal toxicity in mice by protein purification (Fig. 1B) and toxicity tests. A large oligomeric protein was subsequently identified, and N-terminal sequence confirmed that the protein isolated was Perivitellin-2 (PmPV2) [Pma_3499_0.54 and Pma_3499_0.31 (Sun et al., 2019)].

Purified PmPV2 proved to cause the same neurological effects than previously reported for the whole *P. maculata* PVF, with a LD50,96h of 0.25 mg/Kg after i.p. injection. This pointed out that PmPV2 is responsible of the poisonous effect of snail eggs.

### Structural features of PmPV2

Native PmPV2 is a ∼162 kDa oligomeric glycoprotein that with an anionic detergent separate into a single band of ca. 98 kDa, which upon reduction dissociates into a heavy chain (PmPV2-67) and a light chain (PmPV2-31) (Fig. 1C, D, S1). The heterodimer is joined by a single disulfide bridge between Cys161 (PmPV2-31) and Cys398 (PmPV2-67) as determined by mass spectrometry (Fig. 2A, Table S1). PmPV2-67 has two glycoforms of pI 5.22 and 5.38 while PmPV2-31 has a single form (pI 8.16) as determined in two-dimensional electrophoresis (2DE) (Fig. S1).

**Figure 2.**
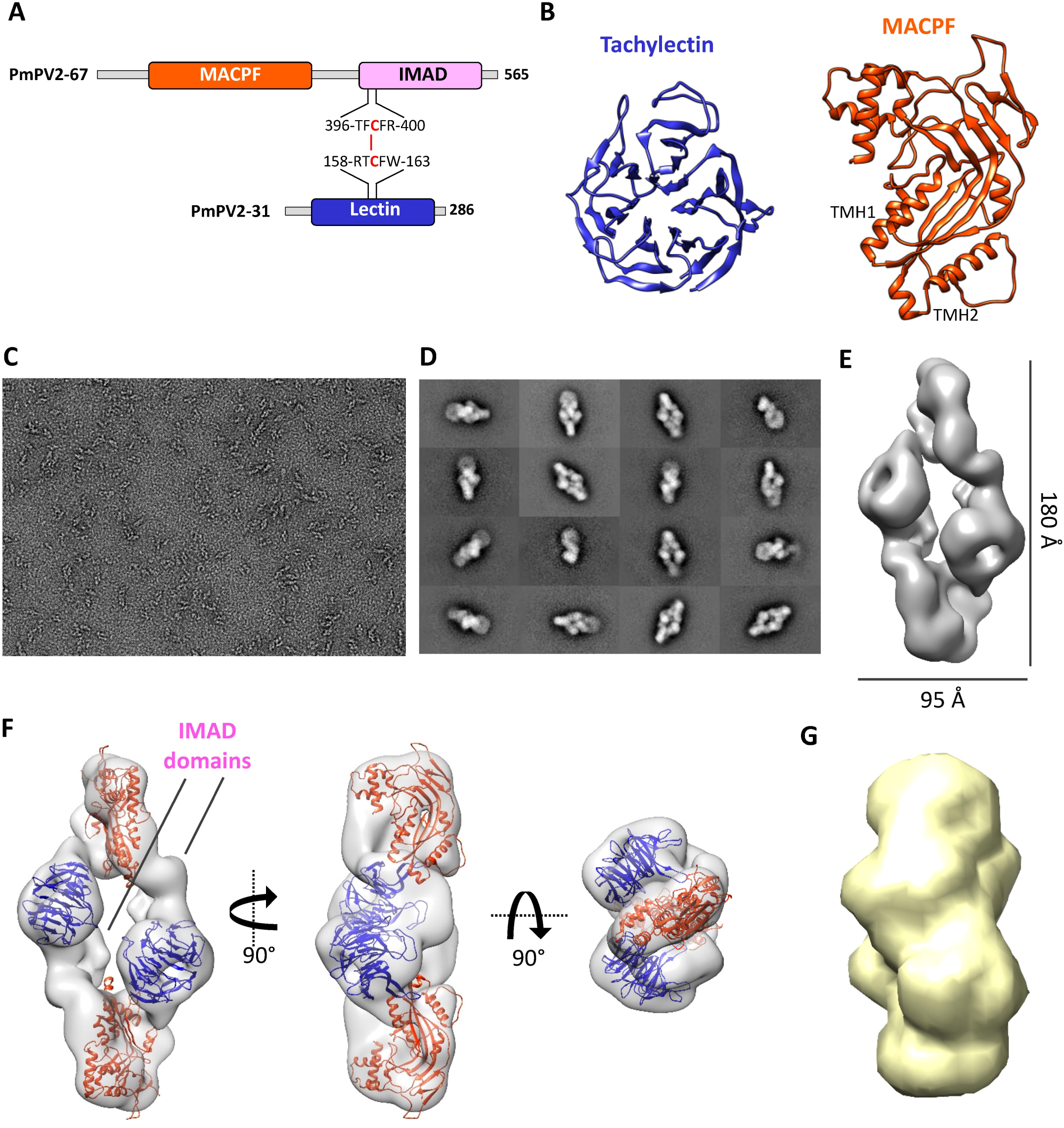
Tertiary and quaternary structure of PmPV2. (A) Schematic architecture of PmPV2. Domains are shown in orange (MACPF) purple (IMAD) and blue (Lectin) boxes. The cysteine residues involved in the interchain disulfide bond are highlighted in red. (B) 3D homology modeling of PmPV2 subunits highlighting characteristic regions of 6-blade β-propeller lectin domain in PmPV2-31 subunit (left), and MACPF domain in PmPV2-67 (right). (C) Representative EM-NS micrograph of PmPV2. (D) Gallery of representative 2D class averages showing the most populated views of the protein. (E) 3D EM map of PmPV2 obtained from reference-free 2D class averages. The monomers (A and B) form AB dimers, which further assemble in a “head-to-tail configuration” fashion as a tetramer. Scale bars are displayed. Rigid-body fitting of MACPF (orange) and Lectin (blue) domains into the NS-EM density map (transparent gray). Different orientation are shown to illustrate the fitted domains across the dimer-of-heterodimers. The models were rigidly docked into the map using UCSF-Chimera. (G) PmPV2 *ab-initio* volume obtained by SAXS (yellow).

Homology modeling of the two subunits allowed us to obtain structures with a reasonable match to their templates (Fig. 2B, Fig S2). Pfam analysis indicates that PmPV2-31 chain has a lectin domain that belongs to the HydWA family (PF06462, E-value=6.6e^-5^) (Fig. S3). This lectin-like domain was identified as structurally similar to carp fish egg lectin (4RUSD), giving a 6 bladed β-propeller structure model (Fig. 2B).

The Nt region of the PmPV2-67 chain has a MACPF domain (PF01823, E-value= 7.e^-24^) with the conserved signature, the Cys residues and the 3 GlyGly sites, all assumed to be important for MACPFs membrane binding (Fig. S3). The best suitable template for the MACPF module was the perforin-1 (3NSJA), an innate immune system protein. The structure modeled was a MACPF fold with their characteristic twisted and bent β-sheet core and its two flanking transmembrane hairpin helixes (TMH1/2) of 40 and 42 residues, respectively (Fig. 2B and Fig. S3). These two helix-clusters are amphiphilic, a known requirement to unfold and insert into membranes.

Although both PmPV2 subunits showed low identity with their templates (Fig. S2), this is in agreement with previous reports for both lectin and MACPF families (ROSADO 2007, SLADE 2008, CHAUDI 2008, KOPEC 2013).

Spectroscopic measurements provided further insight into the structure, indicating that PmPV2 does not have an absorbing prosthetic group (Fig. S4A). Protein tryptophan and other aromatic amino acids are buried in a non-aqueous and highly rigid environment (Fig. S4B, C). The secondary structure, with equivalent amounts of alpha helixes and beta sheets (Fig. S4D and Table S2), agrees with the predicted 3D model.

#### Negative stain reconstruction of PmPV2

We used negative stain electron microscopy (NS-EM) to determine the oligomeric state and obtain low-resolution structural information. Single particles of PmPV2 were perfectly distinguishable (Fig. 2C) from which a preliminary 3D map was obtained.

The ensuing 2D class averages (Fig. 2D) showcased a set of distinct well-defined projections revealing clear structural features. Consequently, an *ab-initio* 3D EM map of PmPV2 was obtained from reference-free 2D class averages, which was iteratively refined imposing a C2 symmetry. The final 3D map (Fig 2E) at 15.2 Å resolution according to 0.5 FSC criteria (Fig. S5), showed size overall dimensions (180 Å x 95 Å), and volume (387.84 Å^3^) consistent with a tetrameric assembly of PmPV2 subunits.

Despite the low-resolution, the tachylectin subunits with the typical donut-like shape (Fig 2 B,E,F), as well as the MACPF subunits with the characteristic planar structure (Fig 2B.E,F), were perfectly recognizable revealing a head-to-tail quaternary rearrangement (Fig 2E).

Accordingly, the docking of the MACPF and tachylectin models into the EM-map, shows an antiparallel dimer-of-heterodimers assembly with a C2 symmetry (Fig. 2E). Both heterodimers are docked to each other by non-covalent forces between a tachylectin from one heterodimer and the MACPF of the other. The rest of the chain does not seem to be part of the tetramer assembly. Despite the PmPV2-67 Ct (IMAD, see below) linker domain is defined in the EM-map between the MACPF and tachylectin domains, no model information is available (Fig 2F).

As a whole, structural data indicate that PmPV2 can be regarded as a dimer of heterodimers held together head-to-tail by non-covalent forces, being the subunits of each heterodimer linked by a single interchain disulfide bond.

#### SAXS of PmPV2

To analyze PmPV2 overall shape and size in solution, we used small angle X-ray scattering (SAXS). Several independent *ab initio* runs yielded reproducible molecular shapes, and the average models generated are consistent with a dimeric state, in agreement with EM-NS and SEC-SLS results (Fig. 2G). SAXS analysis indicate PmPV2 has a gyration radius of 43.9 ± 0.3 Å and a globular and anisometric shape (pair distance distribution *P(r)* with a maximum at 43.7 Å and a *D*_*max*_ of 142.5 Å) compatible with a 173 kDa particle, thus both SLS and SAXS yielded similar molecular weights. The global shape of the low-resolution 3D model of PmPV2 obtained by this method is in good concordance with the NS-EM model (Fig. 2F,G).

### PmPV2 is an active lectin and is able to form transmembrane pores

To analyze the activity of the tachylectin module we tested PmPV2 agglutinating capacity against rabbit red blood cells (RBC). PmPV2 above 0.8 mg/mL produced hemagglutination of intact RBC and also of those pretreated with neuraminidase (Fig. 3A). To demonstrate that the lectin activity was responsible for the agglutination, and not some other process, a control test was performed adding different sugars to inhibit agglutination. This competition assay showed that PmPV2 hemagglutinating activity was strongly inhibited by aminated monosaccharaides, while other sugars had little or no effect (Fig. 3A).

**Figure 3.**
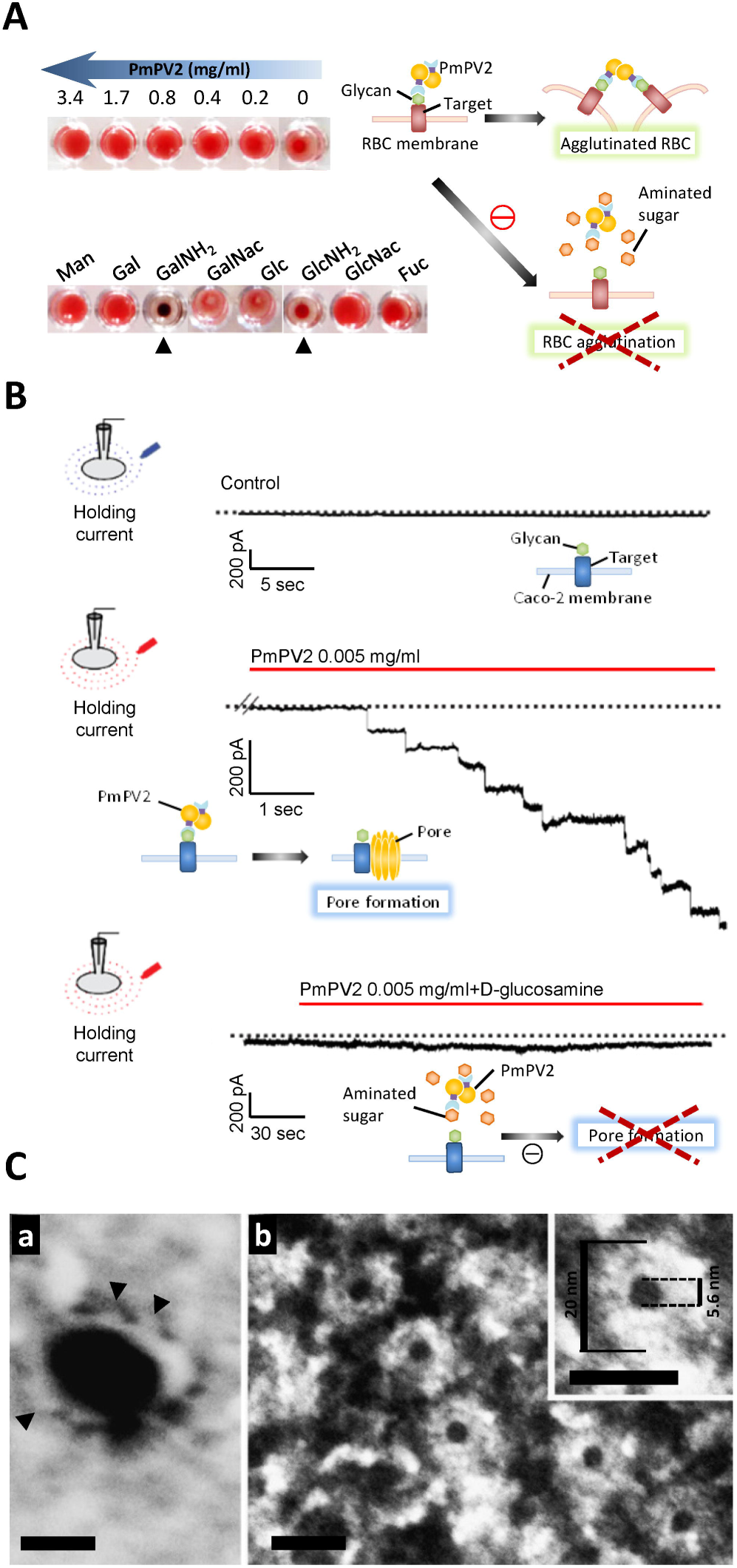
PmPV2 form pores and perforates membranes. (A) Lectin activity of PmPV2 on erythrocytes (upper panel) and hemagglutinating activity of PmPV2 preincubated with monosaccharides (lower panel). D-mannose (Man), D-galactose (Gal), D-galactosamine (GalNH2), N-acetyl-D-galactosamine (GalNac), D-glucose (Glc), D-glucosamine (GlcNH2), N-acetyl-D-glucosamine (GlcNac), L-Fucose (Fuc). (B) Patch clamp experiments: Typical whole cell holding current obtained from a Caco-2 cell continuously clamped at −50 mV, before (upper panel) and after extracellular perfusion of PmPV2 (middle panel) or perfused with PmPV2 preincubated with GlnNH2 (lower panel). (C) TEM imaging of PmPV2 pore formed on liposomes. (a) POPC:Cho liposomes carrying PmPV2 pore-like structures in side view (arrowheads). 50kx amplification. Bar 100 nm. (b) Top view of ring-like structure form by PmPV2 on the liposome surface at 225kx amplification. *Inset:* 640kx amplification. Bar 20 nm.

Considering the presence of a MACPF domain in the PmPV2-67 chain, we also evaluated the putative pore-forming activity of PV2 using Caco-2 cells, a cell line on which it binds to (Dreon et al., 2013). We examined membrane conductance changes by patch clamp techniques. Cells exposed to 29 nM PmPV2 (5 μg/mL) rapidly showed discrete current increments in a stepwise fashion that began to be detectable 2-3 min after the toxin was added (Fig. 3B and Fig. S6). This behavior lasted a few seconds and then the current stabilized at a final increased value respect to the control condition. From each discrete current jump, we calculated the conductance (G), obtaining a mean value of 1,116 ± 53 pS (n = 43 of six cells tested), and estimated a pore diameter (*d) of* 7.2 nm (*d*= 2√ (Gh/σπ)) assuming a solution conductivity (σ) of 1.6 S. m^-1^ and a membrane thickness (h) of 5 nm. This experiment showed that PmPV2 had the capacity to form pores but did not give information on whether the lectin module was needed to recognize and direct the toxin towards the membrane surface. To test this, the toxin was pre incubated with D-glucosamine before adding to the cells. After this treatment, the toxin was unable to change cell conductance, indicating that the lectin module was required for PmPV2 pore formation (Fig. 3B). This result suggested that PFT module was active and dependent on the presence of an active lectin for activity. In agreement with patch clamp results, TEM imaging of PmPV2 interaction with POPC/Cho liposomes captured pore-like structures with an inner diameter of 5.6 ± 0.16 nm (Fig. 3C).

### Phylogenetic analysis revealed a novel MACPF accessory domain exclusive of invertebrates

BLASTp search of PmPV2-31 chain in NCBI non-redundant database revealed 22 similar sequences, mostly belonging to lectin families. All except one, a fish egg lectin-like protein from *Rhinatrema bivittatum*, belonged to invertebrates (Fig. S7A). Remarkably, the Cys161 involved in the disulfide linking of this subunit to the heavy chain, was only observed in *Pomacea* sequences (Fig. S7B). BLASTp analysis of PmPV2-67 chain showed 37 similar sequences scattered in vertebrates and invertebrates, 32 belonging to the MACPF family (Fig. S8). BLASTp searches of this sequence showed two conserved regions: an Nt-region containing the MACPF domain, and a Ct-region with a few matches with unknown proteins. A domain boundary prediction analysis by ThreaDom (Xue, Xu, Wang, & Zhang, 2013) indicated that PmPV2-67 has a relatively disorganized region between the MACPF domain and the Ct-region, suggesting the subunit is composed by two different domains. Analyzing the two regions separately, residues 1-335 (Nt-PmPV2-67) and 336-565 (Ct-PmPV2-67), revealed 32 matching sequences for the Nt region, all belonging to the MACPF family of vertebrates and invertebrates. Unexpectedly, the Ct region matched 18 sequences exclusive of invertebrates (Fig. 4A), 13 associated with Ct-regions of MACPF-containing proteins and 2 associated to a Notch domain (a domain involved in membrane interaction in vertebrate MACPF proteins) (Fig. 4B). Interestingly, phylogenetic analysis indicates an early diversification of MACPF PmPV2 like proteins in Mollusks (Fig. 4A). Multiple sequence alignment of the Ct region with the matching sequences revealed several conserved residues, in particular many Cys (Fig. S7C). We named this novel domain ^..^Invertebrate MACPF Accessory Domain, IMAD^..^. Notably, *P. canaliculata* and *P. maculata* IMADs contain binding site to the tachylectin chain trough a disulfide bridge, thus allowing the MACPF module attachment to the lectin. Other functions of IMAD in invertebrates remain to be investigated.

**Figure 4.**
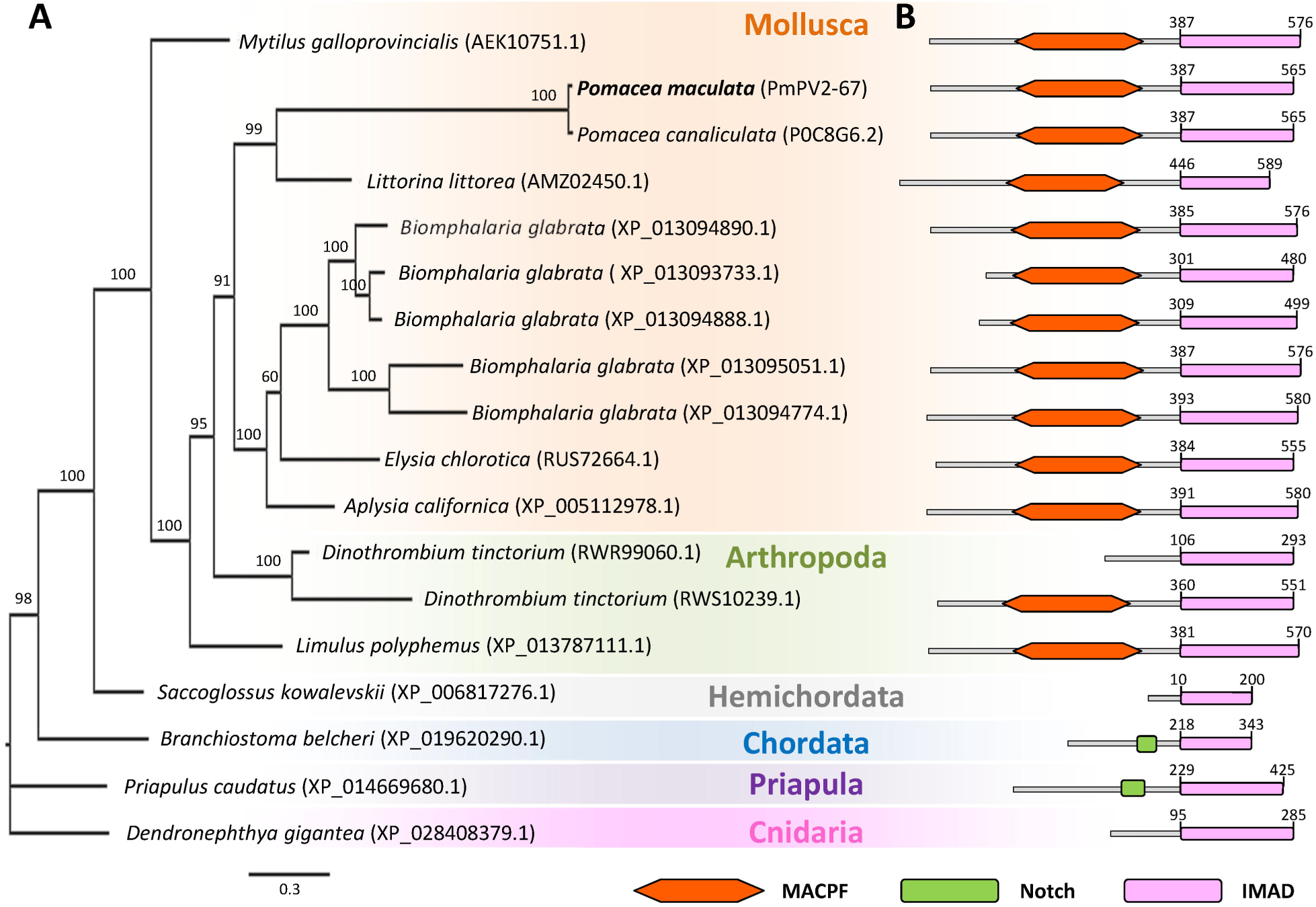
Phylogeny and occurrence of Invertebrate MACPF Accessory Domain (IMAD). (A) Unrooted phylogenetic tree of homologous sequences of Ct-PmPV2-67. Homologues were retrieved from BLASTp analysis, sequences aligned by MUSCLE and phylogeny reconstructed using MrBayes. Node numbers represent Bayesian posterior probabilities (in percentage) of finding a given clade. (B) Domain architecture of Ct-PmPV2-67 homologue sequences highlighting the relative position of the Nt-MACPF and Ct-I MAD domains, as found by ThreaDom and Pfam analysis. AEK10751.1: MACPF domain containing protein *(Mytilus galloprovincialis);* POC8G6.2: Perivitellin-2 67 kDa subunit *(Pomacea canalicu/ata);* AMZ02450.1: Perivitellin-2 67 kDa subunit-like *(Littorina littorea);* XP _013094890.1, XP _013093733.1, XP _013094888.1, XP _013095051.1, and XP _013094774.1: Perivitellin-2 67 kDa subunit-like *(Biomphalaria g/abrata);* RUS72664.1: Hypothetical protein *(Elysia ch/orotica);* XP_005112978.1: Perivitellin-2 67 kDa subunit-like *(Aplysia ca/ifornica);* RWR99060.1, and RWS10239.1: Perivitellin-2 67 kDa subunit-like *(Dinothrombium tinctorium);* XP_013787111.1: Perivitellin-2 67 kDa subunit-like *(Limulus po/yphemus);* XP_006817276.1: Perivitellin-2 67 kDa subunit-like *(Saccog/ossus kowalevskii);* XP_019620290.1: Uncharacterized protein *(Branchiostoma be/cheri);* XP_014669680.1: Uncharacterized protein *(Priapu/us caudatus);* XP_028408379.1: Perivitellin-2 67 kDa subunit-like *(Dendronephthya gigantea)*.

## Discussion

### PmPV2 structure and toxicity

The acquisition of venoms and poisons is a transformative event in the evolution of an animal, because it remodels the predator-prey interaction from a physical to a biochemical battle, enabling animals to prey on, and defend themselves against, much larger animals (Holford, Daly, King, & Norton, 2018). Here we report the initial functional and structural characterization of PmPV2, a toxin that, according to the experimental results on mice and cell cultures, would be a potential defense of apple snail embryos against predation.

Although not as potent as other snail toxins such as conotoxins (Luna-Ramirez et al., 2007), PmPV2 could be consider as “highly toxic”, similar to many snake venoms (Gawade, 2004). The toxin proved lethal to mice when it entered the bloodstream and those receiving sublethal doses displayed neurological signs similar to those caused by the PVF (Giglio et al., 2016) or the PcPV2 toxin (Heras et al., 2008).

The general structural features of PmPV2, analyzed by spectroscopic methods and PAGE, were similar to those previously described for PcPV2 orthologous (Dreon et al., 2013; Frassa, Ceolín, Dreon, & Heras, 2010; Heras et al., 2008). Like PcPV2 (Dreon et al., 2013; Frassa et al., 2010; Heras et al., 2008), PmPV2 sequence indicate the presence of a lectin-like subunit (PmPV2-31) and a MACPF containing subunit (PmPV2-67), sharing 97 % and 96 % similarities, respectively (Mu et al., 2017). In animals, lectins and MACPFs are ubiquitous and typically related with the innate immune system (Anderluh et al., 2014; Rudd, Elliot, Cresswell, Wilson, & Dwek, 2001), the main defense system against pathogens found in invertebrates (Hoffmann, Kafatos, Janeway, & Ezenowitz, 1999). The novelty here is that in PV2s both are combined by a single disulfide bond forming lectin-MACPF heterodimers and two of these heterodimers are held together by non-covalent forces to form the native protein. Therefore, two structural features distinguish the PV2 toxins from the rest of the animal PFTs: (1) they are AB toxins, with a lectin B-chain that binds to cell surface glycans and a MACPF A-chain which kills target cells by forming membrane pores; (2) unlike other MACPF, PV2s are secreted as dimers.

A literature search indicates that PV2s are the only reported animal toxins with a binary AB structure. Furthermore, these are the only AB toxins where the toxic moiety is a member of the MACPF family thus, instead of having toxicity by enzymatic activity to alter target cell metabolism (FALNES 2000), it affects cells by forming pores. From the functional point of view, the lectin in the AB structure would act increasing MACPF targeting specificity as compared to toxins that bind to membranes solely by protein-lipid interactions(Ros & Garcia-Saez, 2015). Remarkably, no dimeric arrangement of AB toxins was reported before.

In our work, SLS, SAXS and NS-EM consistently indicate that native PV2 is a dimer of heterodimers. As far as we know, there is only a single report of another MACPF secreted as a structurally-stable water-soluble dimer (Ellisdon et al., 2015) and not as monomers as the vast majority of MACPF. Interestingly, this dimeric MACPF is also a cytotoxin, the fish stonustoxin (SNTX). However, unlike PV2, SNTX do not have a lectin subunit, or even a carbohydrate-binding domain (Ellisdon et al., 2015). The reason for this dimeric arrangement is still unknown. SAXS and NS-EM derived models allowed a visual analysis of PmPV2, which reveal an antiparallel head-to-tail orientation of its protomers. In the NS-EM 3D reconstruction, the tachylectin subunit appears like a donut, which agreed with the predicted β-propeller structure (Bonnardel et al., 2019; Chen, Chan, & Wang, 2011; Fulop & Jones, 1999; Jawad & Paoli, 2002), whereas MACPF domain presents the characteristic flattened shape of the MACPF/CDC fold involved in oligomerization and pore formation (Rosado et al., 2007). Another interesting aspect of PmPV2 structure is the MACPF Ct domain. In vertebrates Cys-rich Ct-accessory domains are commonly located next to MACPF domains, functioning as ancillary domains key to the MACPF-membrane interaction (Peraro & van der Goot, 2016). Bioinformatic analyses of the PmPV2-67 subunit revealed that in apple snails the MACPF domain is fused with a novel Ct accessory domain, which is likely enhancing its selectivity, membrane binding affinity and/or toxicity (Peraro & van der Goot, 2016; Reboul, Whisstock, & Dunstone, 2016). We found this Ct domain is conserved among many invertebrate MACPF-containing proteins, and phylogenetic analysis suggests that this combination of a MACPF and Ct-domains may have been present in the last common ancestor of invertebrates. We thus propose that this conserved domain, we dubbed IMAD, is a new family of MACPF-accessory domains exclusive of invertebrates with a still unknown structure and a putative membrane recognition function. In *Pomacea*, IMAD is also the binding site to the tachylectin chain. The interaction with several membrane components to attain higher binding affinity and specificity for the target cell has been reported for other MACPF/CDC PFT (Reboul et al., 2016). This is another avenue of future research.

### PmPV2 is a pore-forming toxin (PFT) delivered by a lectin

The presence of a MACPF domain in the primary structure of both PcPV2 and PmPV2 (DREON 2013, MU 2017), suggested a putative pore-forming activity. Here, we confirmed for the first time that PV2s are indeed PFTs and that upon binding, oligomerize into a complex that penetrate the target membrane. Patch clamp experiments also indicated that, once cells are perforated, the membrane oligomeric structures are stable. Besides, the discrete jumps in membrane conductance in a stepwise fashion is consistent with the pore-forming activity already reported for other PFTs (Marchioretto, Podobnik, Dalla Serra, & Anderluh, 2013) (Podack, Ding-E Young, & Cohn, 1986). This was further supported by the identification in the predicted structure of amphipathic sequences in the TMH1/2 together with the typical MACPF/CDC fold required to form pores. Finally, the TEM images provided a visual confirmation of pore-like structures of ∼6 nm inner diameter, which agrees with the pore size estimated by patch clamp measurements, and lies within the range reported for other MACPFs (Anderluh et al., 2014).

We demonstrated that, beside the pore forming activity, PmPV2 is also an active lectin with a primary specificity for aminated sugars. In this regard, CBMs in other AB toxins are found to function in delivering the toxic component of the protein to cell surfaces through glycan-CBM interactions (Boraston, Lammerts van Bueren, Ficko-Blean, & Abbott, 2007). As blocking the lectin activity inhibited the pore-forming capacity on biological membranes, we could suggest that the binding of the tachylectin subunit is a necessary step for the pore formation by the MACPF chain. However, further studies are needed to unveil the membrane binding and pore-formation mechanisms of this toxin.

### Ecological and evolutionary implications

We found that apple snail eggs have evolved a novel PFT, which, combined with other defenses of the egg, would disable essential physiological systems in prey. The toxin shows no resemblance with other gastropod toxins such as echotoxin-2 (Kawashima, Nagai, Ishida, Nagashima, & Shiomi, 2003) or conotoxins from Conidae (Olivera, Rivier, Scott, Hillyard, & Cruz, 1991; Olivera, Showers, Watkins, & Fedosov, 2014). The combination of two unrelated polypeptides resulted in a novel protein with toxic properties, a feature not concurring with the roles classically ascribed to either animal lectins or MACPFs; they have co-opted into a new PFT that would function in *Pomacea* embryo defenses against predation. This has proven successful for the snails as virtually no predator has been able to neutralize this toxin so far (Yusa et al., 2000).

Remarkably, co-occurrence of a MACPF and a tachylectin in non-ampullariids is restricted to an amphioxus, a reptile and a snail. However, there is no information regarding their toxicity, or whether they are covalently linked in those organisms. On the contrary, a recent genomic analysis of tachylectin and MACPF genes showed that they comprise a MACPF-tachylectin complex exclusive of the ampullariid family. This cluster went through several tandem duplications in *P. maculata*, with some copies exclusively expressed in the female albumen gland –the gland that synthesize the egg fluid- and detected in the eggs (Sun et al., 2019). These snails have therefore evolved an optimized defense where a genetically encoded toxin that is maternally deposited in the eggs is at the same time a storage protein for the nutrition of the embryos (Heras, Garín, & Pollero, 1998). Furthermore, it has been suggested that the acquisition of toxic PV2s may have enabled terrestrial egg-laying (Sun et al., 2019). A similar dual function has also been recognized in plant seeds where toxic lectins can also double as storage proteins (Lundgren, 2009).

In conclusion, we provide the first evidence that PV2 toxins from snail eggs are active PFTs. Apple snail PV2, however, differs in several respects from known MACPF pore-forming toxins as it is disulfide-linked to a lectin into an AB toxin arrangement and also because it is secreted as a dimer instead of a monomer in aqueous solutions. Linking two immune proteins in a new toxic entity massively accumulated in the eggs is likely to represent the key step for PV2 novel role in defense against predation, an unparalleled example of protein exaptation. To the best of our knowledge, this is the first description of an animal AB toxin directed toward cell membranes. Future work will look at whether there are differences in the pore structure and oligomerization mechanism between PV2s and other PFTs.

## Methods

### Eggs collection and PmPV2 purification

Adult females of *Pomacea maculata* were collected in the Parana River in San Pedro (33°30’35.97’’ S; 59°41’52.86’’ W), Buenos Aires province, Argentina and kept in the laboratory (Collection permit number DI-2018-181-GDEBA-DAPYAMAGP, Government of the Buenos Aires Province). Eggs were collected within 24 h of laid and kept at −20 °C until processed. Pools of three clutches were homogenized in ice-cold 20 mM Tris-HCl, pH 7.4, keeping a 3:1 v/w buffer:sample ratio as previously described (Heras et al., 2008). The crude homogenate was sequentially centrifuged at 10,000 xg for 30 min and at 100,000 xg for 50 min to obtain the egg perivitelline fluid, PVF.

PmPV2 was obtained following the method described for PcPV2 (Pasquevich, Dreon, & Heras, 2014). Briefly, PVF was ultracentrifuged in a NaBr (density = 1.28 g/ml) gradient at 207,000 xg for 22 h at 4 °C. Then, PmPV2 fraction was purified by high performance liquid chromatography (HPLC) using a Mono Q™ 10/100 GL (GE Healthcare Bio-Sciences AB) column using a gradient of NaCl in 20 mM Tris-HCl buffer, pH 8.5; and by size-exclusion chromatography in a Superdex 200 10/300 GL (GE Healthcare Bio-Sciences AB) column. Purity was checked by electrophoresis in 4-20 % polyacrylamide gels.

Protein content was determined either by the method of Lowry (Lowry, Rosenbrough, Farr, & Randall, 1951) using Bovine Serum Albumin (BSA) as standard, or using PmPV2 molar extinction coefficient at 280 nm, *ε*^*280nm*^ (*see below*).

### Toxicity tests

All studies performed with animals were carried out in accordance with the Guide for the Care and Use of Laboratory Animals (Council, 2011) and were approved by the ‘‘Comité Institucional de Cuidado y Uso de Animales de Experimentación’’ of the School of Medicine, UNLP (Assurance No. P08-01-2013). Animals were obtained from the Experimental Animals Laboratory of the School of Veterinary Science, UNLP. Groups of five female BALB/cAnN mice (body weight: 16 ± 1.1 g) were injected intraperitoneally (i.p.) with a single dose of 200 µL of PBS buffer (1.5 mM NaH_2_PO_4_, 8.1 mM Na_2_HPO_4_, 140 mM NaCl, 2.7 mM KCl, pH 7.4) or the same volume of a serial dilution of five concentrations of PmPV2. Median lethal dose (LD50) was determined by a lethality test 96 h after injection, statistical analysis was performed by PROBIT using EPA-Probit analysis program v1.5 statistical software of the US Environmental Protection Agency (US EPA), based on Finney’s method (Finney, 1971).

### Mass determination

Molecular weight of native PmPV2 in solution was determined by light scattering using a Precision Detectors the column, and the chromatographic runs were performed with a buffer containing 20 mM Tris-HCl pH 7.5, 250 mM NaCl under isocratic conditions at a flow rate of 0.4 mL/min at 20 °C. The concentration of the injected sample was 1.35 mg/ml. The MW of each sample was calculated relating its 90° light scattering and refractive index (RI) signals and comparison of this value with the one obtained for BSA (MW 66.5 kDa) as a standard using the software Discovery32. The reported MW values are an average between the values relating RI and UV with scattering.

### Polyacrylamide gel electrophoresis (PAGE)

Native and subunit composition of PmPV2 was determined by PAGE in 4-20% gradient polyacrylamide gels using Mini-Protean II System (Bio Rad Laboratories, Inc., Hercules, CA). Non-native conditions were performed using 0.1% sodium dodecyl sulfate (SDS), 0.5% dithiothreitol (DTT) and β-mercaptoethanol. Low and high molecular weight markers (GE Healthcare Bioscience, Uppsala, Sweden) were run in parallel. Gels were stained using Coomassie Brillant Blue G-250. Glycosylation was detected by PAS staining following the McGuckin and McKenzie (McGuckin & McKenzie, 1958) method modified by Streitz et al (Streitz et al., 2014), using a commercial Schiff reagent (BioPack). Further analysis was performed by two-dimensional electrophoresis gels (2-DE) in an Ettan IPGphor 3 system (GE Healthcare), as previously described (Pasquevich et al., 2014) using 60 μg of PmPV2.

### Spectroscopic analysis

#### Absorbance

Absorption spectra of PmPV2 (0.64 mg/mL in 20 mM Tris-HCl, 150 mM NaCl buffer, pH 7.5) were recorded between 240 and 700 nm. Ten spectra of three independent pools were measured and averaged. Forth-derivative operation was applied to analyze the relative contribution of different aromatic residues (Butler, Smith, & Schenilder, 1970).

The molar extinction coefficient of denatured PmPV2 was experimentally determined by measuring the absorbance at 280 nm of a solution of 720 μg of lyophilized protein in 6 M guanidinium hydrochloride (GnHCl), following equation 1:

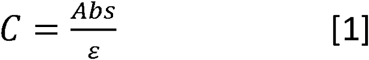

where *C* is the protein concentration (in mg.mL^-1^), *Abs* the absorbance at a given wavelength (in nm), ε the molar extinction coefficient (in mg^-1^.mL). To determine the molar extinction coefficient of the native PmPV2, the absorbance of the native and the denatured protein were measured at identical protein concentrations. Since the concentrations are equal, we combined the equation 1 of the two solutions to obtain the native molar extinction coefficient:

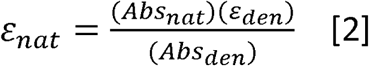

where ε is the molar extinction coefficient (in mg^-1^.mL), *Abs* the absorbance at a given wavelength (in nm), subscript *nat* refers to native protein and subscript *den* refers to denatured protein.

All these experiments were analyzed using an Agilent 8453 UV/Vis diode array spectrophotometer (Agilent Technologies).

#### Fluorescence

Fluorescence emission spectra of PmPV2 (65 µg/mL) in PBS buffer were recorded in scanning mode in a Perkin-Elmer LS55 spectrofluorometer (Norwalk). Protein was exited at 280 nm (4 nm slit) and emission recorded between 275 and 437 nm. Fluorescence measurements were performed in 10 mm optical-path-length quartz-cells. The temperature was controlled at 25±1 °C using a circulating-water bath.

#### Circular dichroism

Spectra of PmPV2 (70–140 μM) were recorded on a Jasco J-810 spectropolarimeter using quartz cylindrical cuvettes of 1-mm or 10-mm path lengths for the far-UV (200–250 nm) and near-UV (250–310 nm) regions, respectively. Data were converted into molar ellipticity [θ]_M_ (deg.cm^2^. dmol^−1^) using a mean residue weight value of 115.5 g/mol for PmPV2.

Proportions of different secondary structures were also obtained using CD spectra in DichroWeb (Whitmore & Wallace, 2008) software using Contin and K2d algorithms.

#### Small angle X-ray scattering (SAXS)

Synchrotron SAXS data from solutions of PmPV2 in 20 mM Tris, pH 7 were collected at the SAXS2 beam line at the Laboratório Nacional de Luz Sincrotron (Campina, Brazil) using MAR 165 CDD detector at a sample-detector distance of 1.511 m and at a wavelength of λ = 0.155 nm (I_(s)_ vs s, where s = 4πsinθ/λ, and 2θ is the scattering angle). Solute concentrations ranging between 0.8 and 2 mg/ml were measured at 20 °C. Five successive 300 second frames were collected. The data were normalized to the intensity of the transmitted beam and radially averaged; the scattering of the solvent-blank was subtracted. The low angle data collected at lower concentration were merged with the highest concentration high angle data to yield the final composite scattering curve, using ATSAS 2.8.4-1 software (Konarev, Volkov, Sokolova, Koch, & Svergun, 2003). *Ab-initio* shape determination was performed using DAMIFF online (https://www.embl-hamburg.de/biosaxs/dammif.html) (Franke & Svergun, 2009) and the resulting damstart.pdb file was used to refine the model using DAMIN (Svergun, 1999) with default parameters. Raw data, fits and models were deposited in SASBDB repository (SASDEN3). (https://www.sasbdb.org/data/SASDEN3/ohuzme8q9a/).

### Determination of disulfide bonds

To identify the disulfide bond between two subunits, 10 μg of purified PmPV2 were first separated by SDS-PAGE and then visualized with colloidal Coomassie Brilliant Blue method. The 98 kDa band was sliced, alkylated with iodacetamide, and digested in-gel with mass spectrometry grade trypsin (Perkin-Elmer).

Peptides were desalted with Sep-Pak C18 cartridges (Waters, Milford, USA) and dried using SpeedVac concentrator (Eppendorf, Hamburg, Germany).Dried samples were reconstituted using 0.1 % formic acid for analysis using LTQ-Orbitrap Elite coupled to an Easy-nLC (Thermo Fisher, Bremen, Germany) with 80 min LC gradient: 5 min in 98% solution A (0.1% formic acid in H2O), 35 min in 7 − 20% solution B (0.1% formic acid in acetonitrile), 20 min in 20 - 35% solution B, 10 min in 35 - 90% solution B, 10 min in 90% solution B. The MS data were captured within a range of 500 to 1800 m/z. The ten most abundant multiple-charged ions with a signal threshold >500 counts were selected for fragmentation under high-energy collision-induced dissociation (HCD; 2.0 m/z of solation width 10 ms of activation time, 40% of normalized collision energy).

Raw data were converted to .mgf files using Proteome Discoverer 1.3.0.339 (Thermo Finnigan, CA). The MS files were searched against custom proteinPmPV2 databases [Pma_3499_0.31, Pma_3499_0.54 and Pma_3499_0.24, which were found in PmPVF (Sun et al., 2019) using the pLink-SS incorporated into pLink 2.3.5 (S. Lu et al., 2018; Shan Lu et al., 2015; Yang et al., 2012) with the cross linkage search of disulfide bond and default parameters.

### Bioinformatic analysis

PmPV2-31 and PmPV2-67 subunit sequences were annotated as Pma_3499_0.54 and Pma_3499_0.31, respectively, by Sun et al. (Sun et al., 2019). PmPV2 related-sequences from different organisms were retrieved from NCBI non-redundant database by BLASTp set as default (threshold E-value=1e-5) and aligned using MUSCLE multiple alignment tool (https://www.ebi.ac.uk/Tools/msa/muscle/) for homology analysis. Phylogenetic analysis was performed using MrBayes v.3.2.6 software, with four chains of 100,000 generations. The tree was sampled every 100 generations, and the final burnin value was set to 20,000. The standard deviation of the split frequencies fell below 0.05. Trees were visualized by FigTree v.1.4.3.

Three-dimensional structures of PmPV2 subunits were predicted by homology modeling using Phyre2 software, which applied a profile-profile alignment algorithm (Kelley, Mezulis, Yates, Wass, & Sternberg, 2015), and pdb files were visualized using UCSF Chimera 1.14 (Pettersen et al., 2004). Quality of the predicted structures was evaluated using NT-PROCHECK software. Models have a confidence level of 100 % in Phyre2, and have more than 90 % residues in the most favored and additional allowed regions in PROCHECK analysis.

### NS data acquisition of PmPV2, image processing, single-particle reconstruction, and refinement

PmPV2 protein samples were suspended in buffer 20 mM Tris-HCl, 150 mM NaCl, pH 8.5 at 0.05 mg/ml and kept on ice before grid preparation (higher concentrations caused oligomerizarion of the samples on the grids). Then, 3 µl of sample was loaded on ultrathin holey-carbon-supported grids, previously pretreated with a glow discharge system for TEM grids during 50 s, under a pressure of 37 Pa. The samples were incubated with the grids 1 min, blotted by filter papers, and then stained with uranyl acetate 2% (w/v) for 30 s. The excess of stain was removed by blotting. PmPV2 EM analysis were performed at LNNano-CNPEM, Brazil (proposal ID 24346). Data acquisition was performed using a Talos F200C (Thermo Fisher) operated at 200 KV with a FEI BM-Ceta direct electron detector model. Data acquisition was performed on a grid, using at a nominal magnification of 73,000 X, corresponding to a calibrated pixel size of 2.02 Å per pixel and a defocus range of −2.0 to −4.0 μm. A total number of 60 micrographs were recorded with an average electron dose per image of 20 e- per Å2. Estimation of CTF, particle picking, 2D classification, reconstruction of an *ab-initio* model, and refinement were executed using the software cisTEM (Grant, Rohou, & Grigorieff, 2018). Briefly, after estimating CTF, an initial template-free particle picking was performed. The preliminary set of picked single particles (20,115 particles) was first exposed to an initial 2D classification resulting in 19 classes (15,317 particles). Subsequently, 2D class averages were used for getting a preliminary *ab-initio* 3D map which was used as a reference for the refinement iterative cycles against 15,317 particles applying a C2 symmetry. The estimated average map resolution was 15.2 Å (FSC=0.5).

The final EM map was sharpened with the auto sharpen tool (Terwilliger, Sobolev, Afonine, & Adams, 2018) from PHENIX (Adams et al., 2010). The models were manually adjusted as rigid bodies using UCSF CHIMERA (Pettersen et al., 2004). After fitting of the models in one half of the dimeric complex, the other half was then independently fitted into the density map. Figures were generated using UCSF-CHIMERA. The NS-EM datasets are available in the wwPDB repository (https://deposit.wwpdb.org/deposition/) with accession code EMD-21097.

### Lectin activity of PmPV2

Rabbit blood samples were obtained from animal facilities at Universidad Nacional de La Plata by cardiac puncture and collected in sterile Alsever’s solution (100 mM glucose, 20 mM NaCl, and 30 mM sodium citrate, pH 7.2). Prior to use, red blood cells (RBC) were washed by centrifugation at 1,500 xg for 10 min in PBS. Hemagglutinating and hemolytic activity were assayed using a two-fold serial dilution of PmPV2 (3.4 mg/mL) as previously described (Dreon et al., 2013). Primary specificity was determined by a competition assay. Erythrocytes were incubated with PmPV2 (0.87 mg/mL) in the presence of 0.1 M of D-mannose, D-galactose, D-galactosamine, N-acetyl-D-galactosamine, D-glucose, D-glucosamine, N-acetyl-D-glucosamine or D-fucose. PmPV2 concentration was selected as the concentration providing visible agglutination in previous analysis. All monosaccharides were purchased from Sigma-Aldrich.

### Microscopic analyses of PmPV2 pores

#### Preparation of small unilamellar vesicles (SUVs)

Multilamellar vesicles were prepared by mixing synthetic 1-palmitoyl-2-oleoyl-sn-glycero-3-phosphocholine (POPC) and cholesterol (Cho) (Avanti Polar Lipids, Birmingham, AL, USA) dissolved in HPLC-grade chloroform/methanol (3:1 molar ratio). Then samples were dried by evaporating the solvent under a stream of nitrogen and then with high vacuum for 2 h in a speed vac. The samples were hydrated in a desired volume of buffer (25 mM HEPES, 150 mM NaCl, pH 7.4) with stirring to facilitate dispersion. Multilayered vesicles were sonicated in an FB-15049 sonicator bath (Fisher Scientific Inc., Waltham, MA, USA) at 30 °C for 1 h to obtain SUVs for AFM and TEM experiments.

#### Transmission electron microscopy (TEM) imaging

An excess of SUVs was incubated with 2.1 μM PmPV2 for 1 h at 37 °C. After treatment, 50 μL of the liposome suspension was placed onto a 300-square-mesh copper grid covered with a Formvar carbon support film (Micro to Nano VOF, Netherlands) and fixed for 1 min. Samples were then negative stained with 50 μL of a 1% (w/v) phosphotungstic acid solution for 30 s. Images at different amplifications were taken using a TEM/STEM FEI Talos F200X microscope (Thermo Scientific) at 200 keV.

### Patch-clamp recordings

Caco-2 cells were allowed to settle onto the cover glass bottom of a 3 ml experimental chamber. The cells were observed with a mechanically stabilized, inverted microscope (Telaval 3, Carl Zeiss, Jena, Germany) equipped with a 40x objective lens. The chamber was perfused for 15 min, at 1 ml.min-1 by gravity, with extracellular saline solution before the patch-clamp experiment was started. Application of test solutions was performed through a multi barreled pipette positioned close to the cell being investigated. All experiments were performed at 22 °C. The standard tight-seal whole-cell configuration of the patch-clamp technique was used (Hamill, Marty, Neher, Sakmann, & Sigworth, 1981) following two different protocols. First, the cells were clamped using voltage ramps from - 50 mV to +60 mV and the macroscopic evoked currents were measured before and after adding PmPV2 to a final concentration of 0.05 mg/mL and after washing cells with the extracellular solution. Secondly, cells were clamped at a holding potential of −50 mV, hence evoking a macroscopic holding current, which we measured before and after adding PmPV2 to a final concentration of 0.005 mg/mL to the bath solution. Glass pipettes were drawn from WPI PG52165-4 glass on a two-stage vertical micropipette puller (PP-83, Narishige Scientific Instrument Laboratories, Tokyo, Japan) and pipette resistance ranged from 2 to 4 MOhms. Ionic currents were measured with an Axopatch 200A amplifier (Axon Instruments, Foster City, CA) filtered at 2 kHz, and digitized (Digidata 1440 Axon Instruments, Foster City, CA) at a sample frequency of 20 kHz. The extracellular saline solution used for recording whole cell ionic currents had a composition similar to the physiological extracellular solution containing 130 mM NaCl, 4.7 mM KCl, 2.5 mM CaCl2, 6 mM glucose, and 5 mM HEPES; the intracellular solution had 130 mM KCl, 5 mM Na2ATP, 1 mM MgCl2, 0.1 mM EGTA, and 5 mM HEPES. The pH of both solutions was adjusted to 7.4 and 7.2, respectively, with NaOH

### Accession numbers

The NS-EM density map has been deposited in the EMBD under accession code EMD-21097. The SAXS data has been deposited in the SASBDB under accession code SASDEN3.

### Availability of data and material

The datasets generated during the SAXS experiments are available in the SASBDB repository (https://www.sasbdb.org/data/SASDEN3/ohuzme8q9a/). The NS-EM datasets are available in the wwPDB repository (https://deposit.wwpdb.org/deposition/) with accession code EMD-21097. All other data generated or analysed during this study are included in this published article and its supplementary information files.

## Acknowledgements

SI, LHO, JC, VM and HH are members of CONICET, Argentina. MSD is member of CIC.BA, Argentina. TB and MLG are Doctoral and Postdoctoral students, respectively with scholarships from CONICET. We thank L. Bauzá for her technical assistance. We are grateful to Single Particle Cryo-EM staff from Brazilian Nanotechnology National Laboratory (LNNano, Campinas) for providing access to their facilities and for the technical assistance during EM experiments. We thank LNLS - Brazilian Synchrotron Light Laboratory for access to their facilities and for the technical assistance during SAXS experiments.

## Declarations

### Ethics approval and consent to participate

The experiment with mice was approved by the “Comité Institucional para el Cuidado y Uso de Animales de Laboratorio” (CICUAL) of the School of Medicine, Universidad Nacional de La Plata (UNLP) (Assurance No. P 01012016) and were carried out in accordance with the Guide for the Care and Use of Laboratory Animals (Guide for care and use of laboratory animals. Washington: Academic Press; 2011).

### Authors’ contributions

SI MG MSD SM VM HH conceived and designed the experiments. MG SI JC SM TB VM MSD JI LHO performed the experiments. MG JI JWQ MSD SI VM SB EP JC LHO HH analysed the data. HH JWQ JC VM LHO contributed reagents/materials/analysis tools. MG SI MSD JC SM TB LHO VM JWQ JI HH wrote the paper.

