## Supplementary material for "Exaptation of two ancient immune proteins into a new dimeric pore-forming toxin in snails": Suplementary files

### Supplementary Information

**Table of Contents**

**A)** Supplementary Figures and Legends……………………………………………………………..1

Figure S1: Structural features of PmPV2.

Figure S2: Homology models of PmPV2 superimposed with crystals.

Figure S3: Deduced amino acid sequences of PmPV2 subunits.

Figure S4: Spectroscopic characterization of PmPV2.

Figure S5: Fourier shell correlation (FSC) curve for the 3D density

Figure S6: Patch clamp of Caco-2 cells.

Figure S7: Multiple alignment of PmPV2 subunits.

Figure S8. Phylogeny of PmpV2 subunits

**B)** Supplementary Tables………………………………………………………………………………………10

Table S1: Inter-chain disulfide bond identification?

Table S2: PmPV2 secondary structure analysis from CD spectra.

### A) Supplementary figures

#### Supplementary Figure 1

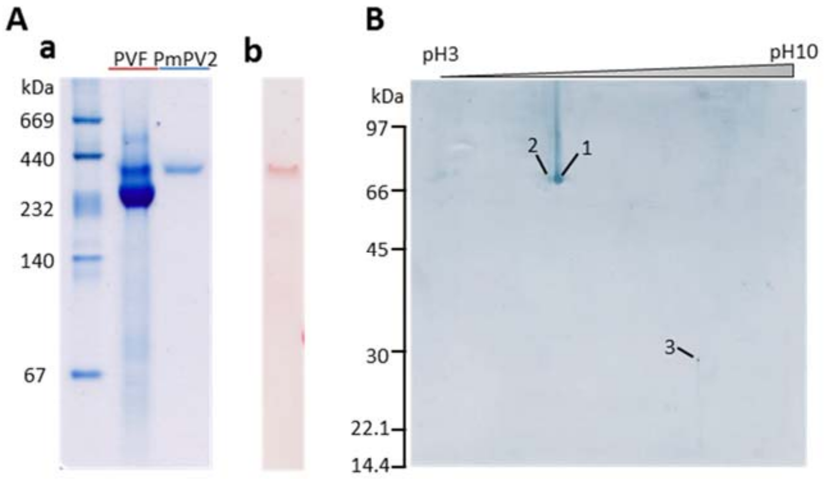


**Figure S1. Structural features of PmPV2. A:** Native polyacrylamide gel electrophoresis (PAGE) in gradient 4-20% stained with Coomassie G-250 (a) or PAS (b). PVF: perivitelline fluid. **B:** Two-dimension electrophoresis; pH ranged from 3.0 to 10.0. Second dimension is a 12 % SDS-PAGE. Spots 1-2: PmPV2-67 kDa subunit isoforms (coordinates 68 kDa, pI=5.22 and 68 kDa, pI 5.38); Spot 3: PmPV2-31 kDa subunit (coordinates 30 kDa, pI 8.16).

**Supplementary Figure 2**

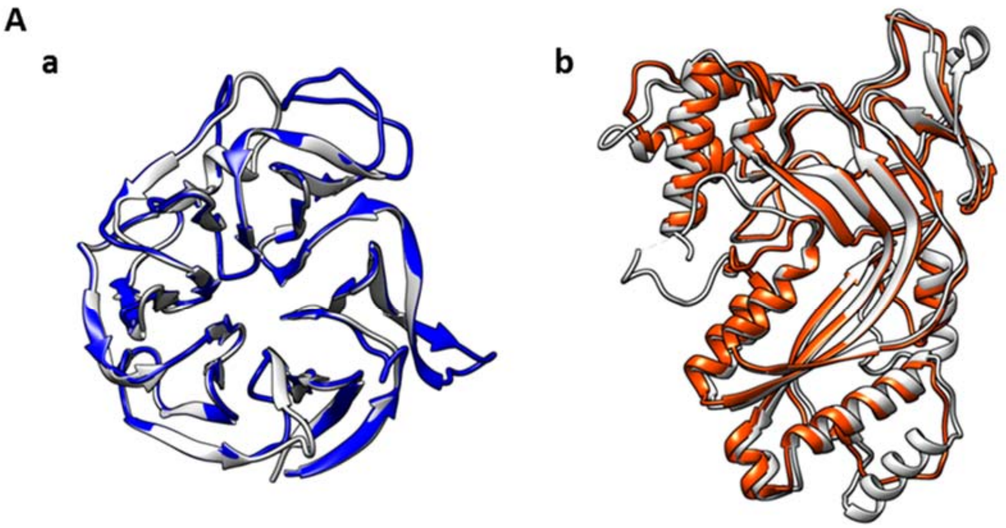


**B**

| Parameter | PmPV2-31 | PmPV2-67 |
|---|---|---|
| % identity* | 22 | 17 |
| RMSD*[a] | 0.8 | 1.3 |
| Z-score*[a] | 39.7 | 38.7 |
| % confidence*[b] | 100 | 100 |
| % most favored + allowed regions[c] | 95.3 | 96.7 |

**Figure S2. Homology modeling of PmPV2 superimposed with cristals. A:** (a) structural superimposition of the PmPV2-31 model (blue) and the fish-egg lectin (grey) (b) structural superimposition of the perforn-1 domain of Nt-PmPV2-67 model (orange) and the perforin-1 (grey). PmPV2 models were obtained using the Phyre2 server. Fish-egg lectin (4RUSD) and perforin-1 (3NSJA) were the best suitable templates found for PmPV-31 and Nt-PmPV2-67, respectively. Structure superposition were obtained using SuperPose server and visualized with USFC Chimera X. No suitable template was found for the Ct region of PmPV2-67 (IMAD) and was therefore excluded in this analysis. **B:** Table comparing  PmPV2 subunits and its templates and model accuracy evaluation. RMSD: root mean square deviation.

* comparison between the model and its corresponding template.

[a] parameters determined by DALI server for structure comparison.

[b] parameter determined by Phyre2 server for homology modeling.

[c] parameter determined by PROCHECK tool for stereochemical quality evaluation.

**Supplementary Figure 3**

**PmPV2-67 [Pma_3499_0.31]**

```
MSQLRWWVVS QVLLLIAICS LDHSEGARVC PKIVPGLDKL RVGVDITKLD LLPLFDLGDN 60
GFRSAVADYT CDRGQTAVVD GESFDVPDQV DSVVIESSGQ QTSSVTTIKS ESQISQALSI 120
SAGISVETAK AGFSSSASYA EMQEAITKYG RTVSQMSAVY TTCSANLSPN LLLGQNPLQT 180
LSRLPSDFTA DTQGYYDFIK TYGTHYFNKG KLGGMFLFTS ETDMSYFQNK NSQQIEATVK 240
ATFASILSTE TGGSSDESKE VIEFKESSLI TSKFFGGQTN LAADGLTKWQ PTIAKLPYFM 300
SGTLSTISSL IADTTKRASM ELAVKNYLLK AKVANLDRLT YIRLNSWSVG HNELRDLSAQ 360
LQNLKTKTIF SDADEKLLQS IEDQVSVPAW FSDRTTFCFR STAVGSADQC NGQSTNTLCA 420
EPNRYTQQYM DKTYLGDTGC RLVWKISTTE STDWFKSVKV NFRWYPTWSP CACGPVGTPF 480
TISAPANSWT QDYLDVTNPK FGECMLQWMI EVPPTATLWA KNLEFCIDFT CGKKKQCVDA 540
NQWTEPYLDI SAHEACGMSW ALIAK 565
```

**PmPV2-31 [Pma_3499_0.54]**

```
MVKKIHFVME RHASIVAFLL AVLALTESQA FTSVKLPRDE HWPYNYVSVG PAGVWAVNRQ 60
NKLFYRTGTY GDNANMGSGW QFKQDGVGQV DVGKDKVGYI NLSGGSLFRI EGISQANPVG 120
GTPKSWEWWT KYIGMSLRED TRFSSRIENQ NKVLTFTFRT CFWASRITNW CFADSSYTET 180
VTAGGSGTWI TKSQLKYKSG TFGNPDTEGG DWILVDSGSF QHVSSGSGVV LAVRSNGELV 240
QRTGITCSLP QGTGWTSMLN SMSRVDTYGT VAWAVDTAGD LYFINL 286
```

**Figure S3. Deduced amino acid sequences of PmPV2 subunits.** Signal peptides are highlighted in green. MACPF domain is in orange, Invertebrate MACPF Accessory Domain (IMAD) is in pink, and HydWA (lectin) domain is in blue. Transmembrane helix 1 (TMH1) is underline in orange and TMH2 in red. Cysteines intervening in the interchain disulfide bond are highlighted in red and underlined. Modified from Sun et al., 2019 doi: https://doi.org/10.1093/molbev/msz084

**Supplementary Figure 4**

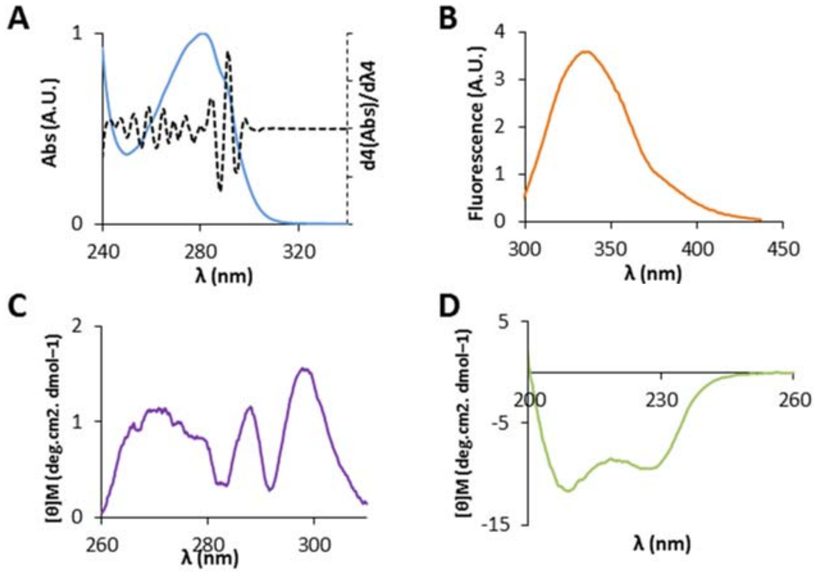


**Figure S4. Spectroscopic characterization of PmPV2. A:** Absorption spectra of PmPV2 (full line), with its fourth derivative spectra (dashed line). **B:** Fluorescence spectrum at 25 °C. **C:** CD spectrum of PmPV2 in the far-UV. **D:** CD spectrum of the PmPV2 in the near-UV region.

**Supplementary Figure 5**

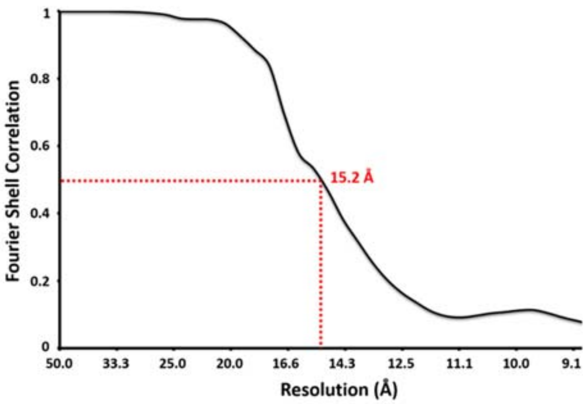


**Figure S5. Fourier shell correlation (FSC) curve for the 3D density map after final iterations of the refinement.** The resolution is estimated to be approximately 15 Å [according to 0.5 FSC criteria] as indicated by the dotted red-line.

**Supplementary Figure 6**

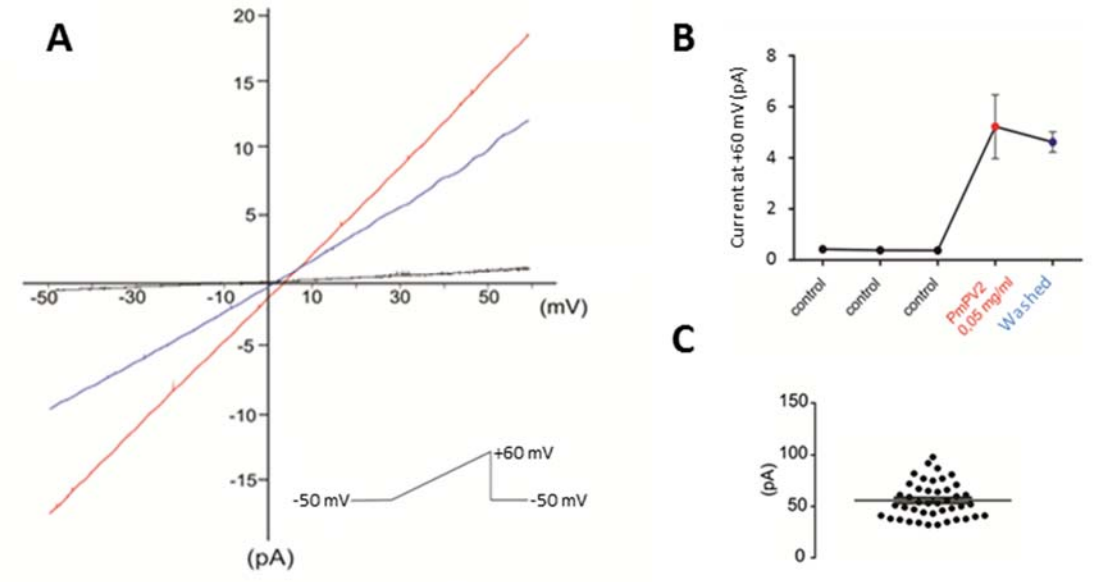


**Figure S6. Patch clamp of Caco-2 cells. A:** typical recording of whole-cell current evoked by a voltage ramp from -50 mV to +60 mV plotted vs. voltage in control conditions (black line), after treatment with PmPV2 (red line) at a final concentration of 0.05 mg/mL, and the same cell washed after treatment with PmPV2 (blue line). **B**: Mean current value measured at +30 mV of control cells (black dots), cells treated with PmPV2 (red dot), and cells washed after treatment with PmPV2 (blue dot). **C:** Current amplitude values of individual current jumps caused by PmPV2 at a final concentration of 0.005 mg/mL.

**Supplementary Figure 7**

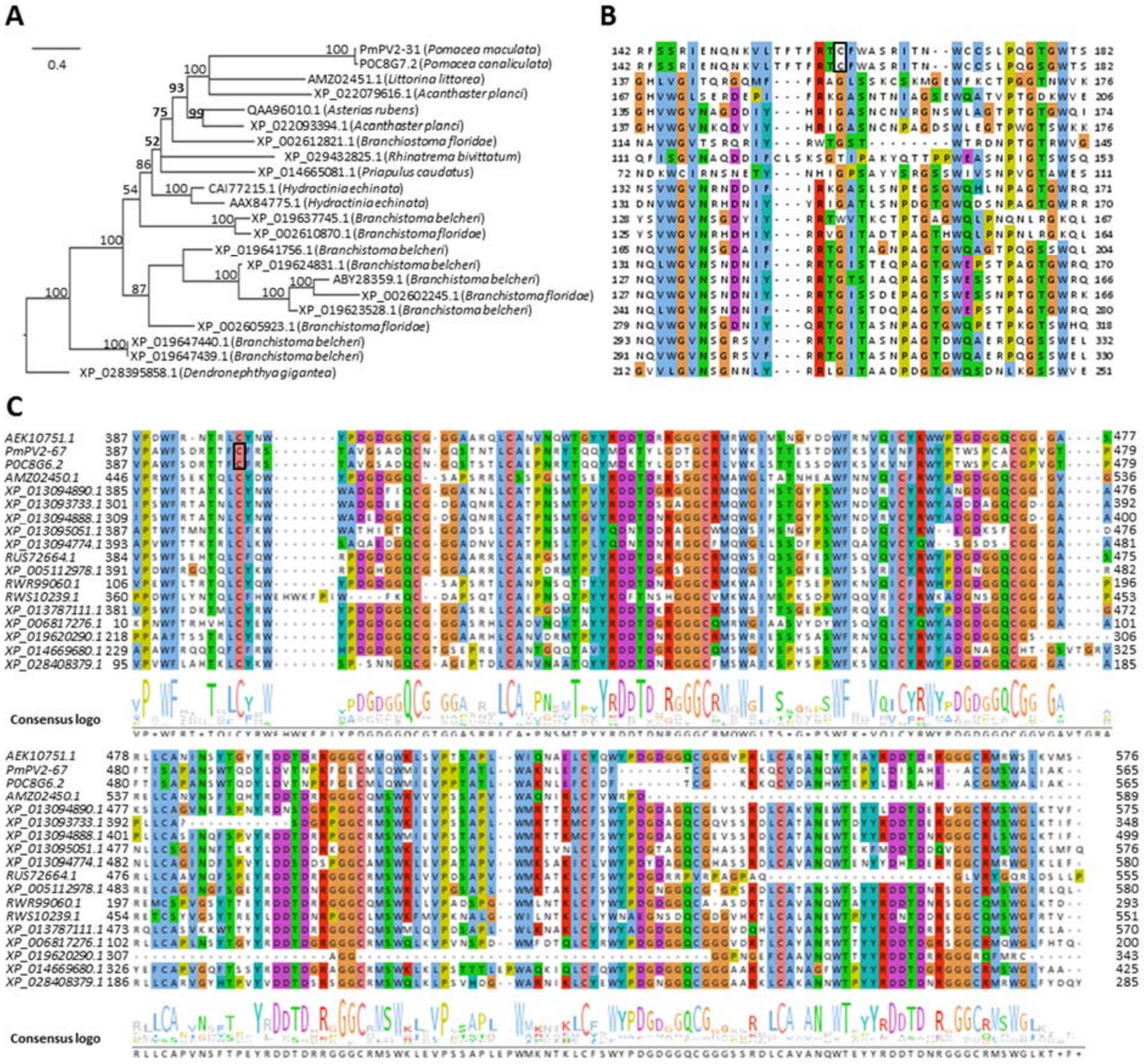


**Figure S7. Comparison between PmPV2 subunits and homolog sequences.**

(A) Unrooted phylogenetic tree of homologous sequences of PmPV2-31. Homologues were retrieved from BLASTp analysis, sequence aligned by MUSCLE and phylogeny reconstructed using MrBayes. Node numbers represent Bayesian posterior probabilities (in percentage) of finding a given clade.

(B) Sequence alignment of PmPV2-31 highlighting the region containing the *Pomacea*-exclusive Cys (boxed) that links the subunit with the IMAD domain.

(C) Sequence alignment of Ct-PmPV2-67 IMAD domain and the consensus logo. *Pomacea* Cys intervening in the interchain disulfide bond is boxed.

Protein sequences for PmPV2-31: P0C8G7.2 : Perivitellin-2 31 kDa subunit (*Pomacea canaliculata*); AMZ02451.1: Perivitellin-2 31 kDa subunit-like (*Littorina littorea*); XP_022079616.1: Tectonin beta-propeller repeat-containing protein 1-like (*Acanthaster planci*); QAA96010.1: Tachylectin-like protein (*Asterias *rubens*); XP_022093394.1: lectin L6-like (*Acanthaster planci*); XP_002612821.1, XP_002610870.1, XP_002602245.1, and XP_002605923.1: Hypothetical protein (*Branchiostoma floridae*); XP_029432825.1: Fish-egg lectin-like (*Rhinatrema bivittatum*); XP_014665081.1: Perivitellin-2 31 kDa subunit-like (*Priapulus caudatus*); CAI77215.1 and AAX84775.1: Tachylectin-like protein (*Hydractinia echinata*); XP_019637745.1: Uncharacterized protein (*Branchiostoma belcheri*); XP_019641756.1: Tectonin beta-propeller repeat-containing protein 2-like (*Branchiostoma belcheri*); XP_019624831.1, and XP_019623528.1 : Tectonin beta-propeller repeat-containing protein 1-like (*Branchiostoma belcheri*); ABY28359.1: Tachylectin-like protein (*Branchiostoma belcheri*); XP_019647440.1: Peroxidasin homolog isoform X2 (*Branchiostoma belcheri*); XP_019647439.1: Peroxidasin homolog isoform X1 (*Branchiostoma belcheri*); XP_028395858.1: Uncharacterized protein (*Dendronephthya gigantea*).

Protein sequences for PmPV2-67: AEK10751.1: MACPF domain containing protein (*Mytilus galloprovincialis*); P0C8G6.2: Perivitelline-2 67 kDa subunit (*Pomacea canaliculata*); AMZ02450.1: Perivitellin-2 67 kDa subunit-like (*Littorina littorea*); XP_013094890.1, XP_013093733.1, XP_013094888.1, XP_013095051.1, and XP_013094774.1: Perivitellin-2 67 kDa subunit-like (*Biomphalaria glabrata*); RUS72664.1: Hypothetical protein (*Elysia chlorotica*); XP_005112978.1: Perivitellin-2 67 kDa subunit-like (*Aplysia californica*); RWR99060.1, and RWS10239.1: Perivitellin-2 67 kDa subunit-like (*Dinothrombium tinctorium*); XP_013787111.1: Perivitellin-2 67 kDa subunit-like (*Limulus polyphemus*); XP_006817276.1: Perivitellin-2 67 kDa subunit-like (*Saccoglossus kowalevskii*); XP_019620290.1: Uncharacterized protein (*Branchiostoma belcheri*); XP_014669680.1: Uncharacterized protein (*Priapulus caudatus*); XP_028408379.1: Perivitellin-2 67 kDa subunit-like (*Dendronephthya gigantea*).

### Suplementary Figure S8

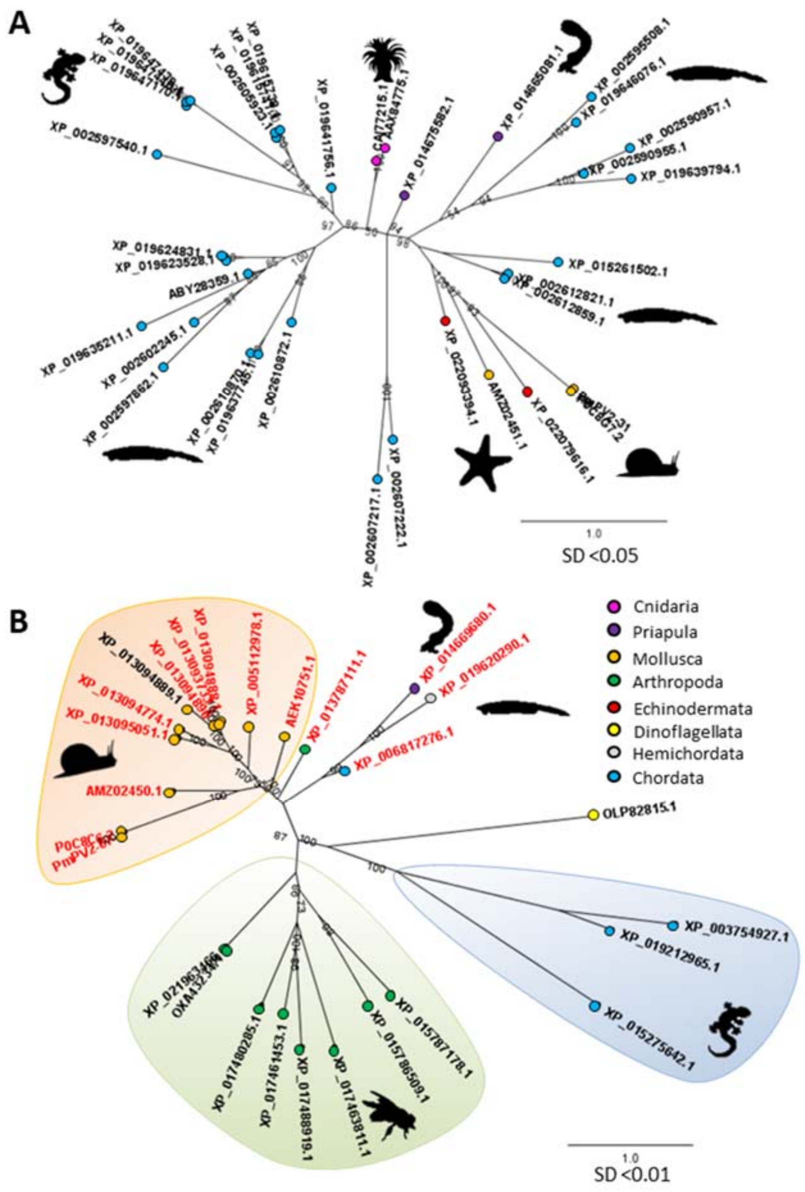


**Figure S8. Phylogeny of PmpV2 subunits. Phylogenetic trees of homologous sequences of PmPV2-31 (A) and PmPV2-67 (B). Phylogeny was generated from sequences obtained by BLASTp using MrBayes and aligned using MUSCLE. SD: mean standard deviation of tree separation. Sequences containing Ct-PmPV2-67 module (IMAD) are highlighted in red. (B) Sequence alignment of PmPV2-31 highlighting the region containing the *Pomacea*-exclusive Cys (boxed) (D) Co-occurrence of MACPF and IMAD domains. (E) Sequence alignment of Ct-PmPV2-67 IMAD domain.**

**Protein sequences for PmPV2-31: P0C8G7.2 *Pomacea canaliculata*; XP_019641756.1, XP_019647440.1, XP_019647439.1 XP_019624831.1, XP_019639794.1, XP_019615747.1, XP_019637745.1, XP_019615738.1, XP_019623528.1, and XP_019646076.1 *Branchiostoma belcheri*; AMZ02451.1 *Littorina littorea*; XP_002612821.1, ABY28359.1, XP_002597540.1, XP_002590957.1, XP_002590955.1, XP_002602245.1, XP_002607222.1, XP_002612859.1, XP_002607217.1, XP_002610870.1, XP_002610872.1, XP_019647170.1, XP_002605923.1, and XP_002597862.1 *Branchiostoma floridae*; XP_022079616.1 and XP_022093394.1 *Acanthaster planci*; CAI77215.1 and**

**AAX84775.1 *Hydractinia echinata*; XP_014675582.1 and XP_014665081.1 *Priapulus caudatus*; XP_015275642.1 *Gekko japonicus*.**

**Protein sequences for PmPV2-67: P0C8G6.2 *Pomacea canaliculata*; XP_013094890.1, XP_013095051.1, XP_013094774.1, XP_013093733.1, XP_013094888.1, and XP_013094889.1 *Biomphalaria glabrata*; AMZ02450.1 *Littorina littorea*; AEK10751.1 *Mytilus galloprovincialis*; XP_005112978.1 *Aplysia californica*; XP_013787111.1 *Limulus polyphemus*; OXA43234.1, and XP_021963466.1 *Folsomia candida*; XP_017480285.1, XP_017463811.1, XP_017488919.1, and XP_017461453.1 *Rhagoletis zephyria*; XP_015786509.1, and XP_015787178.1 *Tetranichus urticae*; XP_014669680.1 *Priapulus caudatus*; XP_006817276.1 *Saccoglossus kowalvskii*; OLP82815.1 *Symbiodinium microadriaticum*; XP_003754927.1 *Sarcophilus harrisii*; XP_019212965.1 *Oreochromis niloticus*; XP_019620290.1 *Branchiostoma belcheri*; XP_015275642.1 *Gekko japonicus*.**

### B) Supplementary Tables

#### Supplementary Table 1

| Identified cross-linked peptides | Number of unique peptide | Number of Spectrum |
|---|---|---|
| Pma_3499_0.31(cys71)-Pma_3499_0.31(cys440) | 1 | 4 |
| Pma_3499_0.31(cys30)-Pma_3499_0.31(cys71) | 1 | 4 |
| Pma_3499_0.31(cys71)-Pma_3499_0.31(cys398) | 1 | 5 |
| Pma_3499_0.31(cys398)-Pma_3499_0.31(cys440) | 1 | 5 |
| **Pma_3499_0.31(cys398)-Pma_3499_0.54(cys161)** | **1** | **5** |
| Pma_3499_0.31(cys398)-Pma_3499_0.31(cys398) | 1 | 3 |

**Table S1. Disulfide bridge on PmPV2 heterodimer.** Cross-linked peptides were mapped to the heavy chain (PmPV2-67) Pma_3499_0.31 and light chain (PmPV2-31) Pma_3499_0.54 with threshold of spectrum >2. Cross-linked peptide of the interchain disulfide bond is highlighted in bold.

**Supplementary Table 2**

| **Algorithm** | **α-helix (%)** | **Β-sheet (%)** | **Disorganized (%)** |
|---|---|---|---|
| K2d | 22 | 23 | 55 |
| Contin | 19,4 | 23,2 | 57,4 |
| **Average** | **20,7** | **23,1** | **56,2** |

**Table S2. PmPV2 secondary structure analysis from CD spectra.** Secondary structure proportions were estimated from CD spectra in the DichroWeb server using the K2d and Contin algorithms.
